# Prolonged reaction times help to eliminate residual errors in visuomotor adaptation

**DOI:** 10.1101/2019.12.26.888941

**Authors:** Lisa Langsdorf, Jana Maresch, Mathias Hegele, Samuel D. McDougle, Raphael Schween

**Affiliations:** NemoLab - Neuromotor Behavior Laboratory, Section Experimental Sensomotorics, Justus Liebig University Giessen, Germany; Center for Mind, Brain and Behavior (CMBB), Universities of Marburg and Giessen, Germany; Department of Brain and Cognitive Sciences, Ben-Gurion University of the Negev, Beersheva, Israel; Department of Psychology, Yale University

**Keywords:** sensorimotor adaptation, reaction time, motor planning, asymptote, explicit strategies

## Abstract

One persistent curiosity in visuomotor adaptation tasks is that participants often do not reach maximal performance. This incomplete asymptote has been explained as a consequence of obligatory computations within the implicit adaptation system, such as an equilibrium between learning and forgetting. A body of recent work has shown that in standard adaptation tasks, cognitive strategies operate alongside implicit learning. We reasoned that incomplete learning in adaptation tasks may primarily reflect a speed-accuracy trade-off on time-consuming motor planning. Across three experiments, we find evidence supporting this hypothesis, showing that hastened motor planning may primarily lead to under-compensation. When an obligatory waiting period was administered before movement start, participants were able to fully counteract imposed perturbations (experiment 1). Inserting the same delay between trials - rather than during movement planning - did not induce full compensation, suggesting that the motor planning interval predicts the learning asymptote (experiment 2). In the last experiment, we asked participants to continuously report their movement intent. We show that emphasizing explicit re-aiming strategies (and concomitantly increasing planning time) also lead to complete asymptotic learning. Findings from all experiments support the hypothesis that incomplete adaptation is, in part, the result of an intrinsic speed-accuracy trade-off, perhaps related to cognitive strategies that require parametric attentional reorienting from the visual target to the goal.

## Introduction

One of the persistent curiosities in studying the human mind is the idea of canonical computations, i.e. that the brain applies similar computations to perform a wide range of different tasks (e.g. Miller, 2016; Pack & Bensmaia, 2015). While most examples for such canonical computations, e.g. linear receptive fields (Movshon et al., 1978,DiCarlo & Johnson, 2000), divisive gain control and normalization (e.g. Carandini & Heeger, 2011) or soft-thresholding of noisy signals (e.g. Ringach & Malone, 2007) have been identified in the fields of neuroscience and artificial intelligence, they have largely eluded scientists in psychology. However, there have been a few but famous instances when psychologists have discovered law-like descriptions of human behavior suggesting the application of similar behavioral algorithms across a wide range of tasks.

One example of such a canonical computation in behavior is the speed-accuracy tradeoff, that is the inverse relation between the accuracy of an action and the time taken to produce it (for a recent review, see Heitz, 2014). The speed-accuracy tradeoff has been shown to shape behavior (a) across domains from motor control (Fitts, 1954; Plamondon & Alimi, 1997) and perception (Grosjean et al., 2007; Ratcliff, 2002) to memory (Hacker, 1980) and mental imagery (Cerritelli et al., 2000; Decety & Jeannerod, 1996), and (b) across species from insects (e.g. Chittka et al., 2003; Ings & Chittka, 2008) and rodents (e.g. Rinberg et al., 2006) to monkeys (Heitz & Schall, 2012) and humans (Wickelgren, 1977).

Another example is the law of practice, according to which performance improvements are generally larger early during practice before they become systematically smaller as practice progresses giving rise to a negatively accelerated relationship between performance and the number of practice trials (Snoddy, 1926, Crossman, 1959, Chen et al., 2005). Regardless of its actual parameters, all versions of the law of practice postulate that performance improvements asymptote, at some point, to a specific performance plateau. For complex skills such as swimming or track and field, it is almost impossible to determine *a priori* the absolute maximum level of performance (but see Berthelot et al., 2015). This is not the case in many experimental paradigms, for instance, in novel visuomotor transformation tasks (e.g., force field adaptation or rotations of visual feedback), where individuals have to adapt an existing movement. These simple manipulations allow for the evaluation of performance improvements relative to an absolute maximum, i.e. an ideal, complete compensation of the transformation (Cunningham,, Lackner & Dizio, 1994, Shadmehr et al.,).

Interestingly, one common observation in this context is that of an incomplete learning asymptote. That is, if individuals are required to make reaching movements and counteract a 30° visuomotor rotation, their performance curve tends to asymptote below full compensation, for instance around ∼25° (Holland et al., 2018, Huberdeau et al., 2015, Haith et al., 2015, Kooij et al., 2016, van der Kooij et al., 2015). This under-compensation leaves a residual performance error significantly different from zero (Hinder et al., 2010, Shmuelof et al., 2012, Spang et al., 2017, Kooij et al., 2016, van der Kooij et al., 2015, Vaswani et al., 2015).

One accepted approach to explain this phenomenon is to leverage state-space models of adaptation, which are incremental Markovian learning algorithms that balance both learning and forgetting during adaptation (Cheng & Sabes, 2006, Smith et al., 2006, Thoroughman & Shadmehr, 2000). When fit to human learning data, most values of learning and forgetting parameters can produce a steady-state equilibrium at an arbitrary asymptote. Consequently, these models provide a natural explanation of the commonly observed undershoot, via an assumption that some amount of forgetting (i.e., reversion to baseline) is inevitable on each trial of the task. This interpretation suggests that incomplete compensation during motor learning is simply a built-in feature of the underlying learning mechanism.

However, Vaswani and colleagues (2015) demonstrated that humans, in principle, possess the capacity to overcome this incomplete asymptote. In their study, visual feedback of their movement was “clamped” after learning; that is the cursor controlled by the participant moved in a fixed trajectory toward the target or to a nearby location with participants only controlling the amplitude. When visual feedback was clamped to a slight deviation from the target with no variability, individuals appeared to adopt a new learning strategy that allowed them to fully compensate a novel visuomotor transformation. To explain this, Vaswani and colleagues (2015) postulated an exploratory learning mechanism that is suppressed by error-based learning. The putative suppressed process only contributes to performance when error-based learning is disengaged, which in their study was caused by a persistent residual error in combination with a contextual change (i.e., the introduction of a lack of natural movement variability) (Shmuelof et al., 2012, Vaswani et al., 2015, Vaswani & Shadmehr, 2013, Wong et al., 2015).

In the present study, we propose and evaluate one alternative, perhaps more parsimonious account of how humans might overcome incomplete asymptotic learning: namely, the level of performance achieved at the later stages of visuomotor adaptation may primarily reflect an intrinsic speed-accuracy tradeoff driven by time-consuming movement planning.

In line with this, research in perceptual decision-making has established that choice reaction time reflects a trade-off between waiting for more information and acting early in order to speed up the accumulation of (uncertain) rewards on future trials (Churchland et al., 2008, Cisek et al., 2009, Thura et al., ; Thura & Cisek, 2017). While visuomotor adaptation tasks traditionally are not studied in the framework of decision-making, recent research has highlighted an important role for volitional decision-making strategies in adaptation tasks (i.e., the explicit re-aiming of movements to counteract perturbations; Bond & Taylor, 2015, Heuer & Hegele, 2015, McDougle et al., 2015, Schween & Hegele, 2017, Taylor et al., 2014). Further evidence suggests that in the context of adaptation to a novel visuomotor rotation such strategies may take the form of mentally rotating the aiming direction of the reaching movement (McDougle & Taylor, 2019), which has been known to require long preparation times (Fernandez-Ruiz et al., 2011, Haith et al., 2015, McDougle & Taylor, 2019). Thus, an incomplete learning asymptote could simply arise from hurried movement initiation leading to prematurely terminating mental rotation of an abstract aiming trajectory during movement planning.

We tested our hypothesis over three behavioral experiments where we artificially extended planning time. We predicted that this simple manipulation would alleviate incomplete asymptotic learning. In the first experiment (experiment 1), we introduced a mandatory waiting period between target presentation and movement onset. In experiment 2, we sought to exclude effects of the total experiment duration by emphasizing the role of within-trial movement planning time versus between-trial consolidation. Finally, in experiment 3, we used an aiming report method (Taylor et al., 2014) to promote the application of explicit motor learning strategies before movement execution and elucidate their influence on the learning asymptote.

## Methods

### Participants

Ninety neurologically healthy and right-handed students from the Justus Liebig University Giessen were recruited as participants (Experiment 1: N = 36, Experiment 2: N = 36, Experiment 3: N = 18) and received monetary compensation or course credit for their participation. Written, informed consent was obtained from all participants before testing. The experimental protocol was approved by the local ethics committee of the Department of Psychology and Sport Science. All participants were self-declared right-handers. Data from one participant (experiment 2) were excluded due to a large number of irregular trials (i.e. premature movement initiation, and moving too fast or too slow).

### Apparatus

Participants sat on a height-adjustable chair facing a 22’’ widescreen LCD monitor (Samsung 2233RZ; display size: 47,3 cm x 29,6 cm; resolution: 1680 × 1050 pixels; frame rate 120 Hz), which was placed on eye level 100 cm in front of them. Their right hand held a digitizing stylus, which they could move across a graphics tablet (Wacom Intuos 4XL). Their hand position recorded from the tip of the stylus was sampled at 130 Hz. Stimulus presentation and movement recording were controlled by a custom build MATLAB script (R2017b), displayed above the table platform, thus preventing direct vision of the hand (left panel Figure 1A).

**Figure 1.**
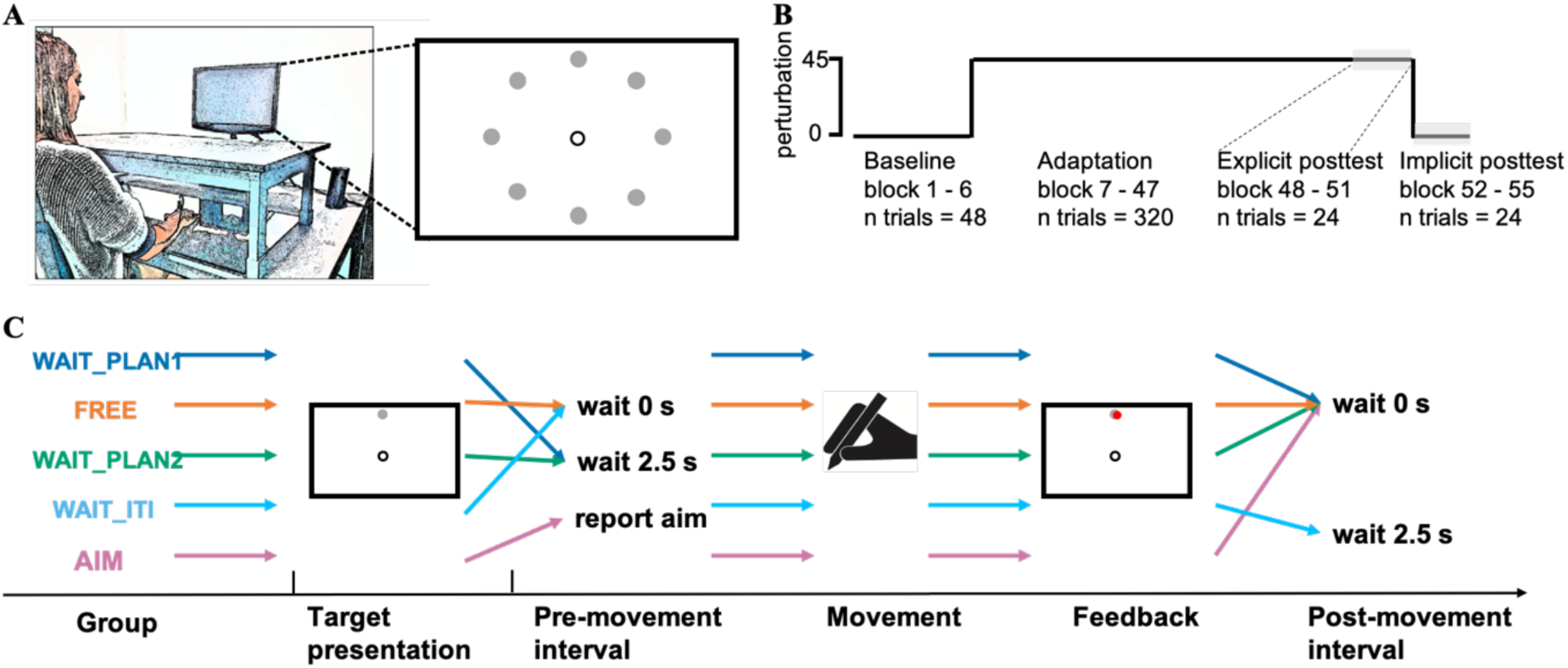
schematic display of the experimental setup (A), overall protocol (B) and sequence of one trial (C). Each participant performed center-out reaching movements with a stylus on the tablet. Visual stimuli and the cursor were presented on a monitor. The visual cursor was displaced according to the protocol (B). During baseline, cursor and stylus position were veridical, during adaptation, the cursor was rotated 45°clockwise relative to the stylus position. Within-trial timing differed between groups (C). Group dependent differences within one trial occurred either during the pre- or post-movement interval. Whereas the FREE and WAIT_ITI groups had no specific task during the pre-movement interval, WAIT_PLAN1 and WAIT_PLAN2 groups were required to wait 2.5 s and the AIM group reported their movement aim. During post-movement interval, only the participants in the WAIT_ITI group were required to wait 2.5 s, whereas all other group continued with the next trial immediately. Panel A adapted from (Schween, Taylor, & Hegele, 2018) under CC-BY-4.0 license.

### Task

Participants performed center-out reaching movements from a common start location to targets in different directions. They were instructed to move the cursor as quickly as possible from the start location in the direction of the displayed target and “shoot through it”. On the monitor, the start location was in the center of the screen, marked by the outline of a circle of 7 mm in diameter. On the table surface, the start location was 20 – 25 cm in front of the participant on the body midline. The target location, marked by a filled green circle of 4 mm in diameter, varied from trial to trial. Targets were placed on an invisible circle with a radius of 100 mm around the start location; target locations were 0, 45, 90, 135, 180, 225, 270, and 315° (0° is from the start location to the right, 90° is forward, 270° is backward; right panel Figure 1A). On baseline and adaptation trials, visual feedback was given by a filled white circle (radius 2.5 mm).

### Design and Procedure

The experiment consisted of three phases: baseline training, training with a 45° clockwise (CW) visuomotor rotation, and posttests (Figure 1B). Baseline training had veridical hand-cursor mapping and was organized into three blocks of eight trials each. In experiment 3, baseline training included three additional blocks in which participants had to report their aiming direction prior to movement onset. Each block consisted of a random permutation of the eight target directions without any direction being repeated in successive trials. Training of the visuomotor rotation of 45° CW consisted of 40 blocks of eight trials each.

The posttest phase consisted of two types of trials: an explicit test (see below) comprising three blocks of eight trials each with each target location occurring once per block, and three blocks of eight aftereffect test trials without visual feedback, with the instruction that the cursor rotation would be absent. In the explicit test trials (Hegele & Heuer, 2010,Heuer & Hegele, 2008), start and target locations were presented together with a white line, centered in the start location with its length corresponding to target distance. Initially, the line was presented at an angle of 180° CCW of the respective target’s direction. Participants instructed the experimenter to adjust the orientation of the line to match the direction of the movement they judged to be correct for the particular target presented.

Each single-movement trial started with the presentation of a white circle in the center of the screen, serving as the starting position for the subsequent reaching movement. In order to help guide participants’ movements back to the start, a white concentric circle appeared after feedback presentation, scaling its radius based on the cursor’s distance from the starting circle. The cursor was displayed when it was within 3 mm of the start location. Once the start position was held for 300 ms, a tone (440Hz, 0.05 ms duration) was presented, followed by a green target (radius 4 mm) appearing in one of the eight target positions and the start circle disappeared. Depending on the assigned group, participants were either instructed to move freely after the target appeared (experiment 1: FREE; experiment 2: WAIT_ITI), to wait 2.5 s for a second tone serving as an imperative (“go” signal) for the movement (experiment 1: WAIT_PLAN1; experiment 2: WAIT_PLAN2), or to report their movement direction and subsequently initiate the reach (experiment 3: AIM).

The white cursor was visible until it exceeded a movement amplitude of 3 mm, after which it disappeared. When the participant’s hand crossed an invisible circle that contained the target, the cursor froze and turned red, providing terminal endpoint feedback for 1.25 s. Movements that fell outside the range of instructed movement time criteria (MT < 100 ms or > 300 ms) were followed by an error message on the screen and the trial was aborted. Those trials were neither repeated nor used in subsequent analyses. If participants moved too soon in one of the waiting groups (before the target appearance or the go cue, see below), they were reminded to wait, and the trial was repeated.

### Groups

The three experiments included five different groups: Two groups of participants took part in experiment 1. One group (N = 19) was instructed to move straight to the target after it appeared with no additional time constraints before moving (FREE). The other group (WAIT_PLAN1, N = 17) was instructed to wait until they heard a high-pitched tone (1000 Hz, 0.05 ms duration) that served as a go-signal. Inspired by previous work indicating that participants are able to mentally rotate their aim 90° off-target within ∼1 s (McDougle & Taylor, 2019), we chose a 2.5 s wait interval to provide ample planning time for the 45° rotation task at hand. The go-signal was presented after this wait interval.

Experiment 2 consisted of two groups: the WAIT_PLAN2 group (N = 22) was a replication of the WAIT_PLAN1 group in experiment 1. Participants in the WAIT_ITI group (N = 20) could initiate movements as soon as the target had appeared on the screen replicating the planning interval of the FREE group from experiment 1. Critcally, the WAIT_ITI experienced an additional 2.5 s waiting period after the presentation of the endpoint feedback. Thus, the two groups, WAIT_PLAN2 and WAIT_ITI, had matched trial lengths but disparate planning intervals. During the 2.5 s inter-trial delay in the WAIT_ITI group, only the target was visible on the screen and participants were told to maintain their final hand position.

Experiment 3 included a single group of participants who were asked to report their aiming direction prior to movement initiation (AIM group, N = 18); (Bond & Taylor, 2015, McDougle et al., 2015,Taylor et al., 2014). The participants in this group saw a numbered ring of visual landmarks. The numbers were arranged at 5.63-degree intervals, with the current target positioned at the “0” position. Clockwise, the numbers became larger, and counterclockwise the numbers became smaller (up to 32, -32, respectively), forming a circle 20 cm in diameter. Participants were instructed to verbally report the number they were aiming their reach at before moving (see (Taylor et al., 2014) for further information on this task). Verbal reports were manually registered by the experimenter on each reporting trial.

## Data Analysis

Position of the stylus on the tablet surface was sampled at 130 Hz and each trial was separately low-pass filtered (fourth-order Butterworth, 10 Hz) using Matlab’s *filtfilt* command, and then numerically differentiated. Tangential velocity was calculated as the Euclidean of x- and y-velocity vectors. Behavior was analyzed in terms of two parameters: reaction time and endpoint error. Reaction time (RT) was calculated as the interval between target presentation and movement onset, which was defined when tangential velocity exceeded 30 mm/s for at least 5 frames (38.5 ms). Endpoint error was calculated as the angular difference between the vector connecting the start circle and the target, and the vector connecting the start circle and the terminal hand position. Endpoint errors were calculated for both training trials and the aftereffect trials. The outcome variable of the explicit perceptual judgement test was calculated as the angular difference between the participant-specified line orientation on the screen and the vector connecting the start and target positions.

For each block of training trials and for the posttest, medians were computed for each participant following screening for outliers. Movements whose endpoint error fell outside three standard deviations of the participants’ individual mean endpoint error in that phase were considered outliers and removed (1.4% of all trials). To compare different levels of asymptote, the last five blocks of the training phase were median averaged and compared between groups using a two-sample Wilcoxon’s rank-sum test. To interpret the effect size, Pearson’s rho and its 95-percent confidence interval was calculated. Statistical analyses were done in Matlab (R2017b) and R (version 3.5.1, http://www.R-project.org/). All results are based on median parametric tests

## Results

### Experiment 1

Experiment 1 tested the speed-accuracy hypothesis by artificially prolonging movement planning time. To do so, we compared two groups. The FREE group could freely initiate their movement, representing a “standard” adaptation experiment. The WAIT_PLAN1 group was required to withhold movement initiation until hearing a “go”-signal 2.5 s after target onset. As shown in Figure 2A, the FREE group displayed the typical incomplete asymptote, whereas the WAIT_PLAN1 group achieved a greater asymptote (mean_WAIT_PLAN1_ = 46.66, sd_WAIT_Plan1_ = 5.85, mean_FREE_ = 41.15, sd_FREE_ = 8.28; V = 244, p = 0.001). Hand directions late during practice were significantly less than 45° in the FREE group (V = 32.5, p = 0.018, r = -0.61, 95% CI = [-0.84, -0.21]), while the WAIT_PLAN1 group did not differ significantly from 45° (V = 108, p = 0.62, r = 0.12, 95% CI = [-0.33, 0.53]).

**Figure 2.**
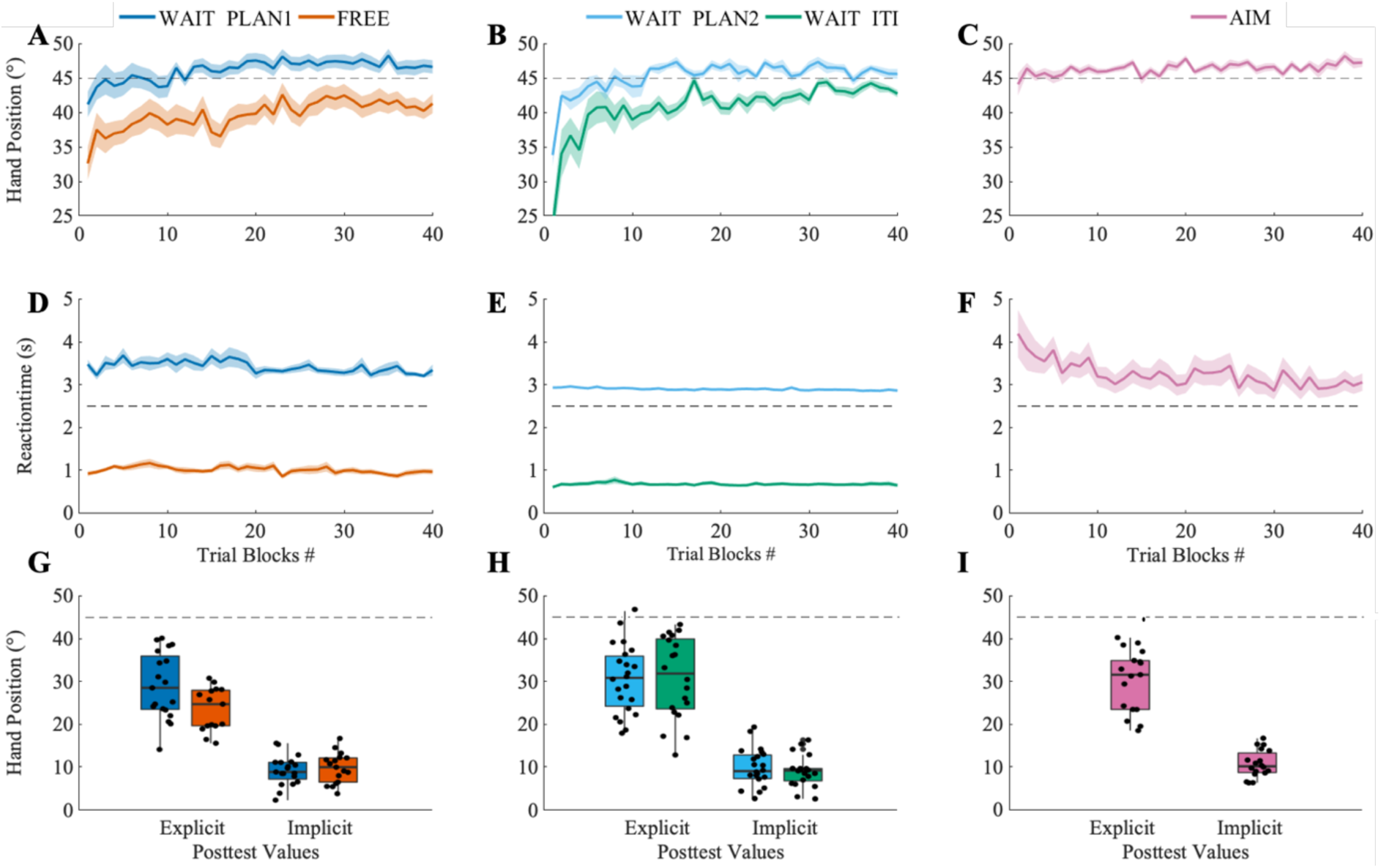
Mean hand direction (panels A-C) and mean reaction times (panels D-F) during practice plotted separately by experiments and groups. Panel G-I show the median hand direction during explicit and implicit posttests, separately and the individual data from single participants. The horizontal dashed lines in panels A-C and H-I indicate ideal compensation for the 45° cursor rotation. In panels D-F, they indicate the forced waiting times of 2.5 seconds in the WAIT_PLAN groups. Shaded error bands represent standard deviation of the mean.

In the explicit judgment test (Figure 2G), the FREE group estimated the rotation to be significantly smaller relative to the WAIT_PLAN1 group (mean_FREE_ = 24.78°, sd_FREE_ = 5.45°, mean_WAIT_PLAN1_ = 30.65°, sd_WAIT_PLAN1_ = 8.33°; V = 81.5, p = 0.036, r = -0.36, 95% CI = [- 0.62, -0.01]). Implicit aftereffects (Figure 2G) did not differ significantly between the groups (mean_FREE_ = 9.99°, sd_FREE_ = 3.81°, mean_WAIT_PLAN1_ = 9.35°, sd_WAIT_PLAN1_ = 3.67°; V = 179, p = 0.59, r = 0.09, 95% CI = [-0.24, 0.39]).

### Experiment 2

Experiment 1 showed that forcing participants to prolong their planning time before movement onset on each trial led to an increase in asymptotic learning. While this observation is consistent with our speed-accuracy trade-off hypothesis, the WAIT_PLAN1 group also exhibited significantly larger amounts of explicit knowledge of the rotation, raising the possibility that this group shows complete asymptote simply because of larger amounts of accumulated explicit knowledge during training. To test this, in experiment 2 we manipulated when the additional waiting time occurred within a trial. If it was a matter of simply building a more elaborate representation of the perturbation by raising awareness and thus accumulating more explicit knowledge of the rotation, then additional processing time between movements should suffice to facilitate complete asymptotic learning. If, on the other hand, the pre-movement planning period was crucial, one would expect that adding time to the interval between the appearance of the target and the signal to initiate the movement would lead to better performance than adding time to the post-feedback interval, i.e. the time interval between the disappearance of terminal endpoint feedback and the onset of the next target. Experiment 2 tested this by contrasting asymptotic learning in a second group that had to wait for 2.5 s during movement planning (WAIT_PLAN2; replication of WAIT_PLAN1) with a group that had to wait for 2.5 s after feedback presentation before the next trial started (WAIT_ITI). In line with our speed-accuracy-hypothesis, inserting waiting time into the planning phase led to an asymptote not significantly different from 45° (V = 235, p = 0.28, r = 0.25, 95% CI = [-0.18, 0.66]) whereas inserting the waiting time into the intertrial interval lead to an asymptote significantly less than 45° (V = 63, p = 0.019, r = -0.44, 95% CI = [-0.75, -0.05]). Those two asymptotes were significantly different from each other (mean_WAIT_PLAN2_ = 46.33, sd_WAIT_PLAN2_ = 3.99; mean_WAIT_ITI_ = 43.96, sd_WAIT_ITI_ = 3.01; W = 311, p = 0.011, r = -0.34, 95% CI = [-0.59, -0.05]) (Figure 2B).

Importantly, for explicit knowledge (Figure 2H), the temporal locus of the additional waiting time did not have a significant effect: Both groups appeared to accumulate equivalent amounts of explicit knowledge (mean_WAIT_ITI_ = 30.53°, sd_WAIT_ITI_ = 8.57°, mean_WAIT_PLAN2_ = 30.88°, sd_WAIT_PLAN2_ = 10.21°; W = 209, p = 0.79, r = -0.04, 95% CI = [- 0.36, 0.25]), but showed greater explicit estimations than the FREE group in experiment 1, whose trial structure did not contain any additional waiting interval (FREE ∼ WAIT_PLAN2: W = 85, p = 0.031, r = -0.37, 95% CI = [-0.63, -0.06]; FREE ∼ WAIT_ITI: W = 93, p = 0.027, r = -0.37, 95% CI = [-0.63, -0.08]). As for implicit aftereffects, both groups in experiment 2 achieved similar results (mean_WAIT_ITI_ = 8.45°, sd_WAIT_ITI_ = 4.77°, mean_WAIT_PLAN2_ = 7.63°, sd_WAIT_PLAN2_ = 3.87°; W = 214, p = 0.89, r = -0.02, 95% CI = [-0.34, 0.36]).

### Experiment 3

In the last experiment, we sought to account for the possibility that it is not time per se, but the increased participation of explicit processes that raises the level of asymptote. We thus instructed participants to verbally report their movement aim prior to movement execution trial-by-trial (Taylor et al., 2014), potentially priming the explicit component of adaptation. We reasoned that this procedure serves as an opportunity to replicate our findings in a procedure that requires active explicit engagement during the planning interval. Compensation for the rotation reached asymptote around 46.63° (sd = 4.12°), which was significantly larger than 45° (V = 125, p = 0.045, r = 0.41, 95% CI = [-0.08, 0.75]), suggesting that adaptation at asymptote was complete and, in fact, overcompensated for the rotation (Figure 2C).

Explicit judgements of required compensation (mean = 28.32, sd = 10.95) (Figure 2I) were significantly less than 45° (V = 0, p < 0.0002, r = -0.88, 95% CI = [-0.88, -0.87]) but significantly greater than 0° (V = 170, p < 0.0002, r = 0.87, 95% CI = [0.82, 0.88]). Implicit aftereffects (mean = 9.38, sd = 3.4) were also significantly different from both 0° and 45° (V = 171, p < 0.0001, V = 0, p < 0.0001, r = -0.87, 95% CI = [-0.88, -0.88], r = 0.87, 95% CI = [0.87, 0.87], respectively). If we assume that the explicit and implicit components are the two main elements in a fully additive model that generates adaptive behavior, the implicit component can be calculated by subtracting the hand position from the aim report (Figure 2L). Comparing those values to the posttest values, we do not find a significant difference, neither in explicit nor in the implicit component (W = 123, p = 0.22; W = 129, p = 0.31, respectively).

To test whether the reporting task influenced the outcome of the explicit judgement tests, we compared the posttest values between the AIM group and those of the other groups in experiments 1 and 2. There was a significant difference in the explicit judgements between the AIM group and the FREE group from experiment 1 (W = 197.5, p = 0.025, r = 0.39, 95% CI = [0.05, 0.6]) but none between the WAIT_PLAN and AIM (W = 160.5, p = 0.76, r = - 0.05, 95% CI = [-0.36, 0.27]). Across the AIM group and WAIT_PLAN2 and WAIT_ITI groups in experiment 2, there were no differences in the explicit judgement tests (W = 160, p = 0.57, r = -0.09, 95% CI = [-0.39, 0.22]; W = 190.5, p = 0.85, r = -0.03, 95% CI = [-0.34, 0.28]). Similar results were observed for the implicit aftereffects: Neither the FREE group, the WAIT_PLAN1 group from experiment 1, nor the WAIT_PLAN2 and WAIT_ITI groups had significantly different aftereffects relative to the AIM group (W = 140.5, p = 0.69, r = -0.07, 95% CI = [-0.38, 0.28]; W = 167.5, p = 0.93, r = -0.01, 95% CI = [-0.35, 0.29]; W = 227.5, p = 0.08, r = 0.19, 95% CI = [-0.13, 0.45]; W = 265.5, p = 0.05, r = 0.25, 95% CI = [-0.07, 0.54], respectively). These results suggest that experimentally querying the explicit process of adaptation does not qualitatively alter the explicit/implicit learning balance but does act to improve the adaptation asymptote by slowing down planning.

## Discussion

This study was designed to investigate whether previously reported findings of incomplete asymptotic visuomotor learning may be reframed, at least in part, as an instantiation of a ubiquitous canonical computation in human information processing: the tradeoff between the speed and accuracy of actions. In line with this hypothesis, artificially prolonging the waiting period prior to the onset of a goal-directed movement elevated asymptotic learning and appeared to eliminate residual errors. This benefit was specific to prolonging motor planning, the time interval between the appearance of the visual target and the go-signal. Prolonging the interval between visual feedback and the start of the next trial (the intertrial interval) did not provide the same benefit to learning. Our results provide support for a parsimonious explanation that time-consuming planning processes are potentially the main driver of incomplete asymptotic learning.

Why did hasty planning result in consistent undershooting rather than both undershooting and overshooting (i.e., greater movement variability)? We propose that parametric mental computations in visuomotor rotation tasks could explain the undershooting phenomenon: In visuomotor rotation tasks, participants’ reaction times increase linearly with the magnitude of the imposed rotation (Georgopoulos & Massey, 1987, McDougle & Taylor, 2019), reflecting a putative mental rotation process (Shepard & Metzler, 1971). Thus, in our framework, undershooting is the consequence of participants not taking the time needed to fully complete a mental rotation of their planned reach trajectory. This view is further supported by the results of our third experiment, in which emphasizing the application of explicit aiming strategies prior to movement initiation led to qualitatively similar asymptotic learning as in the groups with prolonged planning intervals. Note that delaying movement initiation did not only cause full compensation, but induced overcompensation suggesting that implicit processes superimposed onto an accurate explicit rotation strategy may have caused reach angles to drift, gradually adapting the hand further in the direction of compensation (cf. Mazzoni, 2006).

The idea of a speed accuracy tradeoff prematurely interrupting putative mental rotation processes during motor planning also provides an explanation for previously observed age-related differences in visuomotor learning. Hegele & Heuer (2013) used explicit instructions and cognitive pretraining prior to learning a novel visuomotor rotation to boost explicit knowledge of the transformation. Older adults with full explicit knowledge of the transformation turned out to be less efficient in applying it for strategic corrections of their aiming movements. This age-related difference with respect to the behavioral exploitation of explicit knowledge became manifest only when participants had almost perfect explicit knowledge, but not when they had only poor explicit knowledge and thus a small range of associated strategic adjustments at different levels of exploitation. Given the present results, one could speculate that the reduced exploitation of explicit knowledge for strategic corrections in older participants is due to a combination of age-related slowing in mental rotation and the premature termination of (slowed) mentally rotating their aiming direction during motor planning.

Traditionally, the incomplete asymptote phenomenon was explained by state-space models of adaptation (Cheng & Sabes, 2006, Smith et al., 2006, Thoroughman & Shadmehr, 2000), according to which the adapted state reaches an equilibrium between learning from error and decaying towards baseline in each trial. As subsequent studies indicated that this model alone is insufficient for explaining incomplete asymptotic behavior, alternatives were proposed: For example, Vaswani and colleagues (Vaswani et al., 2015) suggested that a process that learns from spatial error feedback suppresses other mechanisms that could drive full compensation (Shmuelof et al., 2012). In our study, participants in all groups received similar spatial error feedback. Thus, a potential suppression should have affected all groups equally, suggesting that spatial error feedback suppressing other learning mechanisms would not be sufficient to explain the modulations in asymptote we observed.

A new approach to the state-space model is that residual errors in adaptation paradigms are caused by implicit processes that tune the sensitivity to errors until it reaches the equilibrium with constant forgetting (Albert et al., 2019). The authors in this recent study manipulated the variability of the perturbation and found that residual errors increase with the perturbations’ variance. We note that, whereas our hypothesis could potentially be adapted to account for these variations in asymptote (e.g. experiencing perturbation variability could affect the benefit that learners expect from planning, and thus the time they spend on it), we did not consider this possibility *a priori* in hypothesis generation. However, we note that in one experiment, this study also showed a speed-accuracy tradeoff by obtaining larger residual errors when the reaction time is artificially shortened compared to free reaction times, regardless of the variance of perturbation. Thus, we argue that additional planning time is an essential element in eliminating residual errors to achieve full compensation, though it need not be the only thing determining the exact asymptotic value.

Moreover, we also note that consistent undershooting relative to the perturbation, as observed here and in previous studies, is critically not seen in experimental paradigms designed to isolate the implicit component of visuomotor adaptation (Morehead et al., 2017)– indeed, even when rotational perturbation are as small as ∼1.75, implicit adaptation appears to asymptote around ∼15° (Kim et al., 2018). These results suggest that claims of an incomplete asymptote within, specifically, the implicit adaptation mechanism must define the asymptote relative to an intrinsic capacity of the system, rather than the size of the visual error. Thus, it may be that incomplete compensation relative to the visual error (i.e., task error) mainly involves cognitive processes like speed-accuracy tradeoffs, as argued here, but incomplete asymptotic performance of the implicit system relative to its own capacity (i.e., responses to sensory prediction error) requires a separate explanation.

Recent accounts have framed motor planning as a time-consuming optimization process from which a reduction in movement accuracy arises naturally when constraints are imposed (Al Borno et al., 2019). Our findings suggest that similar principles apply when one is intentionally choosing to perform a movement in another direction than the one implied by the target presented, and that learners naturally constrain their planning time even in seemingly unconstrained conditions. Haith and colleagues (Haith et al., 2016) recently showed that movement preparation and initiation are independent i.e. that, instead of complete preparation triggering movement initiation, humans appear to determine a time for movement initiation based on when it expects planning to be completed. This view naturally implies the possibility to initiate a movement that has not been sufficiently prepared. The planning time chosen may therefore trade off the accuracy it expects planning to achieve within a given time and an urgency to move on (e.g. fueled by a desire to increase reward rate; Churchland et al., 2008, Cisek et al., 2009, Thura et al.,, Thura & Cisek, 2017).

Many of the common explanations for incomplete asymptote outlined above imply that it is a fundamental property of learning. Psychology and kinesiology traditionally distinguish learning effects from performance effects, where underlying knowledge can be identical in different cases, but retrieval processes in specific test conditions can lead to different performance profiles (Magill & Anderson, 2017, Schmidt & Lee, 2011). Whereas our experiments were not specifically designed to distinguish learning from performance effects, our findings suggest that both may contribute to incomplete asymptote in adaptation. Specifically, explicit knowledge of the rotation magnitude was increased with added planning time in experiment 1, suggesting that some of the benefit of longer planning times may come about by learners honing their explicit knowledge. However, the observation that explicit knowledge was similarly increased regardless of whether additional time was added at the beginning or end of a trial in experiment 2 indicates that this learning effect may be a non-specific consequence of longer ITIs, and that the remaining increase in asymptote is a performance effect. A recent paper analyzing preparatory neural states in rhesus monkeys performing visuomotor learning tasks also found that longer preparation times not only yielded smaller variance on the current trial, but also smaller errors on the subsequent trial, supporting a learning effect (Vyas et al., 2020). Future research could attempt to better delineate learning from performance effects in human motor adaptation.

Lastly, we do not claim that other mechanisms affecting learning do not contribute to asymptotic behavior (Albert et al., 2019), or that a state-space model with gradual decay towards zero is generally invalid (Brennan & Smith, 2015). What we suggest is that one potentially major aspect determining the magnitude of asymptotic errors is a speed accuracy trade-off. Since this decision process is likely to be relevant across a broader range of motor tasks, we speculate that our results extend beyond motor adaptation and that simple interventions, like explicitly prolonging reaction times to allow for complete planning, could improve asymptotic performance in a range of motor learning tasks.

## Author Contribution

LL, JM, MH, SDM and RS conceived and designed research; LL collected data; LL, JM, MH, SDM and RS analyzed data; LL, JM, MH, SDM and RS interpreted results of experiments; LL prepared figures; LL drafted manuscript; LL, JM, MH, SDM and RS edited and revised manuscript; LL, JM, MH, SDM and RS approved final version of manuscript

